# Deep Profiling of Plasma Proteoforms with Engineered Nanoparticles for Top-down Proteomics

**DOI:** 10.1101/2024.07.20.604425

**Authors:** Che-Fan Huang, Michael A. Hollas, Aniel Sanchez, Mrittika Bhattacharya, Giang Ho, Ambika Sundaresan, Michael A. Caldwell, Xiaoyan Zhao, Ryan Benz, Asim Siddiqui, Neil L. Kelleher

**Affiliations:** Proteomics Center of Excellence, Northwestern University, Evanston, IL 60208, USA; Seer Inc., Redwood City, CA 94065, USA

**Keywords:** top-down proteomics, proteoforms, nanoparticles, protein corona, plasma

## Abstract

The dynamic range challenge for detection of proteins and their proteoforms in human plasma has been well documented. Here, we use the nanoparticle protein corona approach to enrich low-abundant proteins selectively and reproducibly from human plasma and use top-down proteomics to quantify differential enrichment for the 2841 detected proteoforms from 114 proteins. Furthermore, nanoparticle enrichment allowed top-down detection of proteoforms between ∼1 µg/mL and ∼10 pg/mL in absolute abundance, providing up to 10^5^-fold increase in proteome depth over neat plasma in which only proteoforms from abundant proteins (>1 µg/mL) were detected. The ability to monitor medium and some low abundant proteoforms through reproducible enrichment significantly extends the applicability of proteoform research by adding depth beyond albumin, immunoglobins and apolipoproteins to uncover many involved in immunity and cell signaling. As proteoforms carry unique information content relative to peptides, this report opens the door to deeper proteoform sequencing in clinical proteomics of disease or aging cohorts.

## INTRODUCTION

Human plasma is an easily accessible, minimally invasive sample type that contains valuable biological information for biomarker discovery and clinical applications.^1–3^ However, analysis of the plasma proteome is challenging due to the overwhelming presence of high-abundance proteins that make up more than 90% of the total plasma proteome, including albumin, immunoglobulins, and other abundant proteins.^4^ Due to the wide dynamic range of protein concentration spanning >11 orders of magnitude, molecular weight (MW) fractionations,^5^ depletion of high-abundant proteins,^6–8^ or enrichment of less abundant proteins^9, 10^ are often required prior to proteomics analysis with mass spectrometry.

To address dynamic range challenge in analyzing plasma proteome, the Proteograph^TM^ workflow was established to facilitate deep and broad plasma proteomics measurement at scale.^11–13^ The workflow includes contacting biofluids with engineered nanoparticles (NPs) to form protein a corona. Varying the physicochemical properties of the nanoparticles results in distinct protein enrichment behaviors, which can be analyzed by mass spectrometry using bottom-up proteomics (BUP) with enzyme digestion. This approach allowed the detection of over 8000 protein groups,^11, 14^ enabling increased depth for plasma proteome profiling and biomarker discovery. For example, it was used to identify distinct BMP1 isoforms in non-small cell lung cancer subjects.^15^ The automatic workflow also has advantages for large scale studies including multiple sample types and/or big cohort analysis. Recent work by Van Eyk and co-workers on high-throughput biomarker discovery across 9 sample types reported that Seer engineered nanoparticles led to reliable quantification of 3359 protein groups using a 24-minute liquid chromatography-mass spectrometry (LC-MS) gradient, which included 137 out of 216 FDA approved circulating biomarkers.^16^ Wilcox and co-workers combined the Proteograph with timsTOF HT technology for deep plasma protein biomarker discovery and reported over 4000 quantifiable protein groups.^17^ Together, these applications show that the protein corona enrichment using Seer NPs and the Proteograph workflow enables robust deep sampling prior to mass spectrometry-based proteomics.

There is a growing interest in proteoform level information acquired by top-down proteomics (TDP) to address the limitations of protein inference and post-translational modification (PTM) analysis in BUP.^18–24^ To link protein composition more directly to function and disease phenotypes, we recently reported that TDP analysis of peripheral blood mononuclear cells (PBMCs) revealed proteoform biomarker candidates for organ transplant rejection that was reproduced in a multi-center trial.^25^ We also demonstrated that TDP captured new differences in patient ApoA-I proteoforms enriched from plasma due to lipidation of Lys88 that correlated better than total ApoA-I to several metrics of cardiovascular disease risk.^26^ Most recently, we showed that proteoforms in plasma can be signatures of liver cirrhosis progression.^27^ These examples indicate that analyzing proteoforms can improve disease diagnosis, understanding underlying mechanisms and identifying treatment targets.

However, TDP of plasma proteoforms also suffers from the wide dynamic range of the plasma proteome and presents the additional challenge that a single gene can give rise to hundreds of proteoforms via combinations of genetic variation, alternatively spliced RNA transcripts, endogenous proteolysis and PTMs. Lee and co-workers used extensive MW fractionation (10-15 <30 kDa fractions) to identify 442 unique proteoforms from 71 proteins.^28^ Fornelli and co-workers reported a record number of 1481 unique proteoforms from 130 proteins using high-field asymmetric waveform ion mobility spectrometry (FAIMS) to further simplify serum proteoform mixtures from only two MW fractions of <30 kDa plasma proteins.^29^ Most recently, Sun and co-workers demonstrated an example of TDP proteoform identification from protein corona.^30^ In their work, polystyrene nanoparticles (PSNPs) were incubated with human plasma to form protein corona, which was then eluted into SDS buffer, cleaned and analyzed with capillary zone electrophoresis (CZE) for identification of 263 unique proteoforms from 50 proteins in the 3-70 kDa size range. While most of the proteoforms came from abundant proteins such as apolipoproteins (ApoA-I-II 100 proteoforms, ApoC-I-III 77 proteoforms), albumin and complement C3, this work hinted the promising future of TDP characterization of protein corona, particularly of those tailored to enrich lower abundant proteoforms.

Here, we present results of TDP analysis of engineered NPs that differentially interrogate human plasma using a modified Proteograph workflow. We observed a record number of unique plasma proteoforms (2841) from 114 proteins in LC-MS with clear evidence of reproducible NP enrichment of plasma proteins. By mapping the identified proteins to the Human Plasma Proteome Project (HPPP) database,^31^ our data showed identification of low abundant proteins at concentrations less than 10 pg/mL by HPPP estimates and associated with pathways in immune responses and cell signaling. This work opens a door for deep quantitative analysis of proteoforms in plasma and could reveal biological differences that were previously hindered by abundant proteins present in plasma.

## EXPERIMENTAL SECTION

### Plasma Sample Preparation

Human plasma samples were purchased from BioIVT where samples were collected from three healthy donors (two Hispanic female, age 35 and 47, and one African American male, age 32). Plasma was used either neat or processed with Seer engineered NPs using a modified Proteograph workflow (XT well A and XT well B). Each well was incubated with 100 µL of plasma and washed as previously described.^11^ For each type of XT well, 40 wells were combined, eluted into 250 µL of SDS lysis buffer in the Proteograph assay kit, and dried in a vacuum centrifuge.

### Molecular Weight Fractionation

Proteins were fractionated using a polyacrylamide-gel-based prefractionation for analysis of intact proteoforms and protein complexes by mass spectrometry (PEPPI-MS)^32^ workflow (<50 kDa) prior to LC-MS analysis. Neat plasma (7 µL, ∼500 µg total protein) and NP eluates (∼250 µg total protein) were resuspended in 40 µL of non-reducing NuPAGE sample buffer (Thermo Fisher Scientific) and boiled at 95 °C for 10 min. Proteins were resolved in a 1.5 mm NuPAGE 4-12% gel (Thermo Fisher Scientific) at 70 V for 10 min and 150 V for 10 min. Bands under 50 kDa were excised, crushed, homogenized, and shaken (1400 rpm, r.t., 20 min) in 100 mM ammonium bicarbonate, pH 9 with 0.1 % SDS. The slurry was transferred to a 0.45 µm centrifugal filter (Corning) and spun at 14,000 rpm for 10 min. The flow-through was stored at -80 °C. Samples were precipitated with methanol/water/chloroform and resuspended in 100 µL (neat plasma) or 50 µL (NPs) sample buffer (94.8% water, 5% acetonitrile and 0.2% formic acid) for LC-MS. A detailed PEPPI-MS protocol can be found in our previous work.^24^

### Top-Down Liquid Chromatography-Mass Spectrometry

Proteins (10 µL each run) were separated using a Vanquish Neo UHPLC chromatographic system (Thermo Fisher Scientific). Reversed-phase LC was performed on a MAbPac EASY-Spray column (150 mm length by 150 µm i.d., Thermo Fisher Scientific) with in-house packed with PLRP-S trap (25 mm length by 150 µm i.d., Agilent). The total run time was 120 min using a gradient of mobile phase A (99.9% water, 0.1% formic acid) and mobile phase B (80% water, 19.9% acetonitrile, and 0.1% formic acid). The flow rate was set at 1 µL/min and the gradient used to resolve proteins was: 5% B at 0 min., 20% B at 5 min, 70% B at 110 min, 99% B from 111 to 114 min, and 5% B from 115 to 120 min. The column outlet was coupled inline to an EasySpray source and an Orbitrap Eclipse mass spectrometer (Thermo Fisher Scientific) operating in intact protein mode with 2 mTorr of N_2_ pressure in the ion routing multiple (IRM). The transfer capillary temperature was set at 320 °C, ion funnel RF was set at 60%, and a 15 V of source CID was applied. MS^1^ spectra were acquired at 120,000 resolving power (at *m/z* 200), normalized AGC target of 1000%, 100 ms of maximum injection time, and 1 μscan. Data-dependent top-N-2sec MS^2^ method used 32 NCE for HCD to generate fragmentation spectra acquired at 60,000 resolving power (at *m/z* 200), with normalized AGC target of 2000%, 1200 ms maximum injection time, and 1 μscan. Precursors were isolated with quadrupole using a 3 *m/z* isolation window, dynamic exclusion of 60 s duration, and threshold of 1 x 10^4^ intensity. All mass spectrometry .raw data and .tdReport files were uploaded to MassIVE (repository number MSV000095086).

### Top-Down Data Search and Analysis

The raw data files were searched with a publicly available TDPortal v4.1.0 (https://portal.nrtdp.northwestern.edu/) workflow based on the Galaxy Project^33^ that generated results reported with a 1% context-dependent false discovery rate (FDR) assigned at the protein, isoform, and proteoform levels.^34^ A proteoform database was created from SWISS-PROT human database (Taxon 9606 – June 2020) that contained 2.4 million proteoform entries. A -14-ppm *m/z* spectral shift was applied to account for instrument calibration. Proteins and proteoforms were filtered for 1% FDR for the identification (upset plots). For quantitative proteoform analysis, a CSV intensity sheet was generated utilizing an isotopic fitting algorithm across the chromatogram to obtain the intensities of all proteoforms identified in the whole study with 10% FDR confidence in each sample. This includes those previously considered as “un-identified” and not meeting the 1% FDR cutoff for identification.^35^ Box-Cox transformed intensity values were subjected to a hierarchical linear model-based ANOVA, with Benjamini and Hochberg FDR correction (α = 0.05), to find proteoforms that were differentially enriched between sample groups. Volcano plots were generated, where each proteoform was represented as a function of estimated effect size (in log_2_ fold-change) and the statistical confidence that differences between the two samples were significantly different (-log_10_ of instantaneous q-values). Q-values above 0.05 were considered significant. Proteoform heatmaps were generated using the ‘ComplexHeatmap’ R package. Identified proteins were matched with the HPPP database^31^ using accession numbers for the waterfall plots. Gene ontology (GO) analyses were performed on the metascape website (https://metascape.org/) v3.5.20240101.^36^

## RESULTS AND DISCUSSION

### Adapting the Proteograph Workflow for Top-Down Proteomics

The Proteograph workflow is well-established for BUP applications. The automatic sample processing steps include protein corona formation on mixtures of NPs engineered to maximize plasma proteome coverage, washes, enzyme digestion and peptide clean-up.^11, 15–17^ Two mixtures of NPs are currently available in the latest generation Proteograph XT workflow: XT well A and XT well B. We modified the workflow with SDS-elution of protein corona for TDP study of plasma from three healthy donors (biorep01-03). As illustrated in the scheme in **Figure 1**, plasma was used neat (7 µL, ∼500 µg total protein) or enriched with XT well A and XT well B (40 wells each) NPs. Then, the proteins were fractionated for <50 kDa (PEPPI-MS). Each sample (including neat and NP-enriched) was prepared in three technical replicates (techrep01-03) and finally analyzed with LC-MS over a 120-min gradient in three injections (injrep01-03). The study involved 81 randomized LC-MS runs.

**Figure 1.**
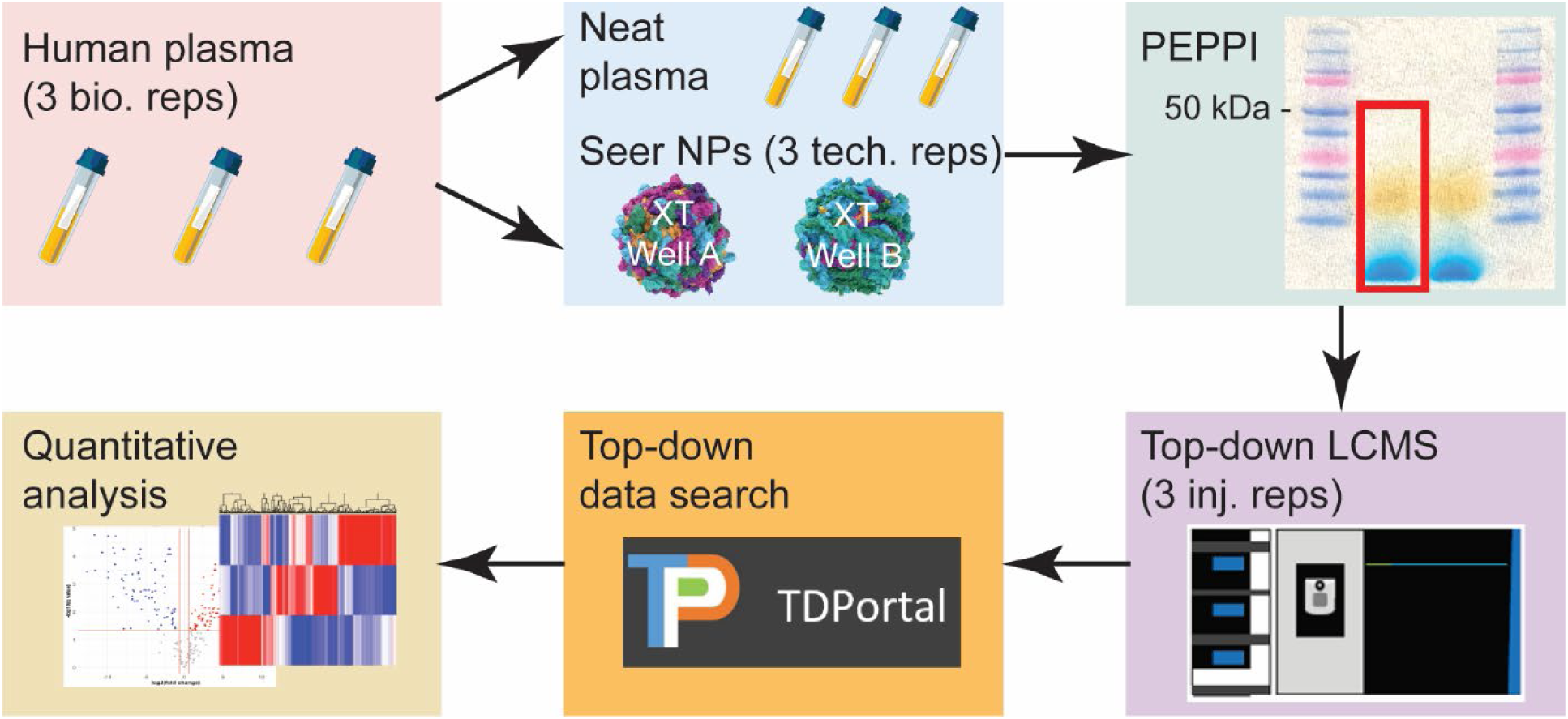
Study design and workflow. Plasma from three human subjects (bioreps) was enriched with two reaction wells from Proteograph XT (XT well A and XT well B). Neat plasma and nanoparticle eluates were extracted for proteins <50 kDa (PEPPI), and an established top-down LC/MS workflow in discovery mode and TDportal search were used to identify and quantify proteoforms. Each nanoparticle enriched sample was performed in triplicate (techreps). All LCMS injections were performed in triplicate (injreps). Proteoforms were filtered for 1% FDR and the quantitative analysis was performed in RStudio.

### Nanoparticles Improve the Coverage of the Intact Plasma Proteome

The data were searched against a curated human proteoform database and hits passing a 1% FDR at the protein and proteoform levels were considered identified (ID). Neat plasma and NPs combined, we identified 2841 unique proteoforms from 114 proteins, a 4-fold and 6-fold increase respectively over neat plasma only conditions. Figure 2A and **2B** show upset plots of proteoform and protein IDs respectively at different intersections of treatments (neat plasma, XT well A and XT well B). Each of the three treatment groups had large numbers of unique proteoforms (673, 676, and 1491 respectively) while the overlaps between neat plasma and NPs were very small. There were only two proteoforms shared between neat plasma and XT well A; 12 proteoforms shared between neat plasma and XT well B; and 13 proteoforms that were identified in all three conditions. Similarly at protein level, there were only 12 proteins shared between neat plasma and NPs. This observation demonstrates that the NP protein corona enrichment enables greater depth in the interrogation of plasma proteome, which is typically hindered by high-abundant proteins in standard TDP and BUP analyses. We also note that there were some overlaps in proteoform (223) and protein (21) IDs between XT well A and XT well B. These two groups of NPs were engineered to interrogate orthogonal groups of plasma proteome while not mutually exclusive as demonstrated by BUP.^11–13^

**Figure 2.**
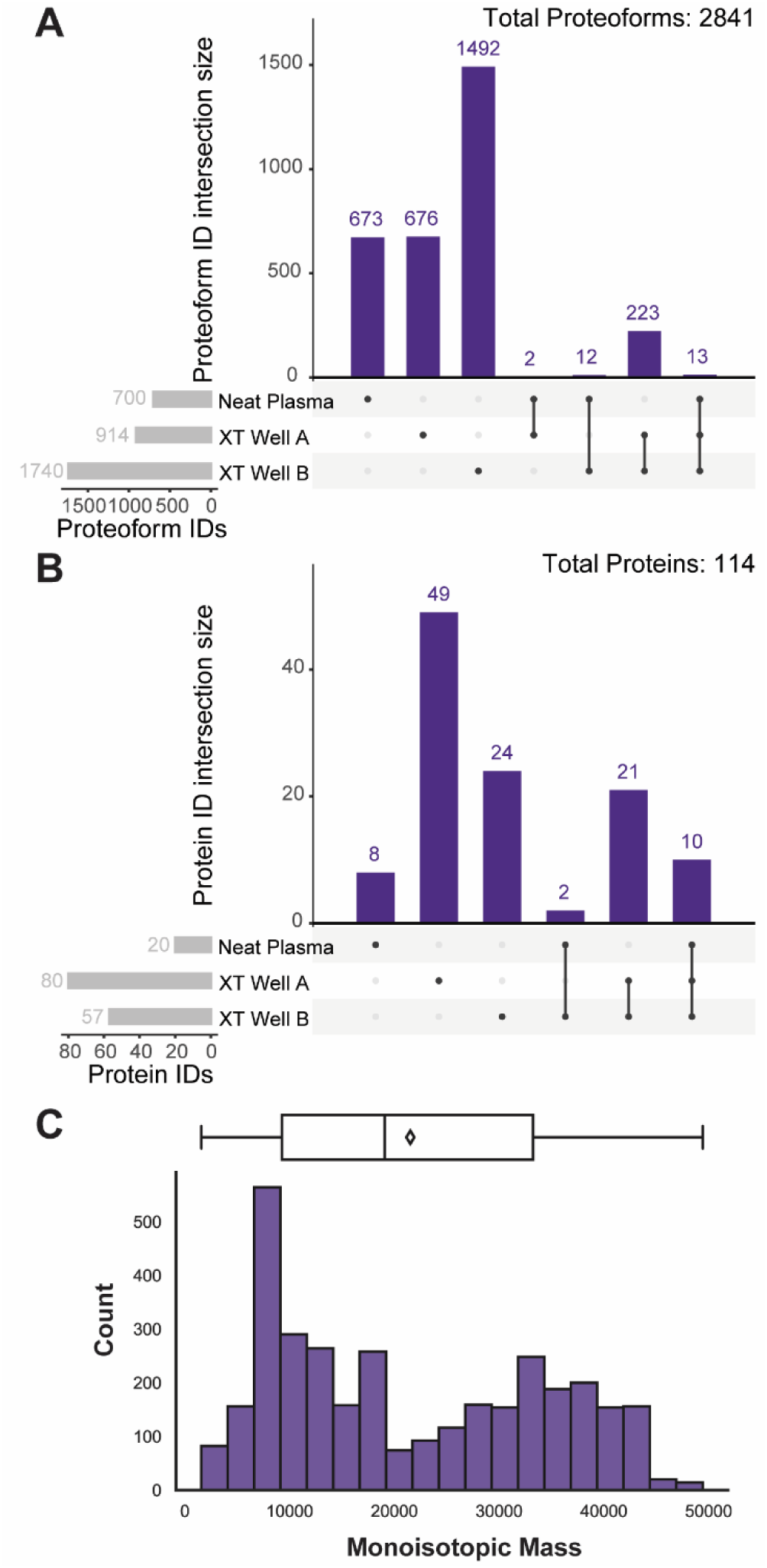
Proteoforms and proteins identified in this study. We report 2841 unique proteoforms (A) and 114 proteins (B) identified in neat plasma and NPs combined. Compared to neat plasma only, the NPs increase the proteoform identifications by over 4-fold. We note that the intersection between neat plasma and XT well A and B are extremely small, supporting that NPs interrogate parts of the plasma proteome that were inaccessible without enrichment. (C) Mass distribution of the identified proteoforms, all of which were <50 kDa using the top-down PEPPI-LCMS workflow.

Figure 2C shows a histogram of proteoform IDs by their monoisotopic mass. The distribution is as expected with predominantly small proteoforms since proteins were specifically sought <50 kDa. However, we note that TDP LC-MS has predominantly been focused on <30 kDa proteoforms in the past due to difficulties in separation, signal dilution in multiple charge states, ion decaying and acquisition of isotopically resolved spectra for identification/quantification.^37, 38^ With the recent improvement in separation technology and MS instrumentation, we can now tackle the proteoforms in 30-50 kDa range with quantifiable data acquired in data-dependent acquisition (DDA) mode. Looking closer into the proteoforms we identified, many of them have canonical sequences above 50 kDa in MW. For example, Prothrombin (accession #P00734) has a canonical sequence of 622 amino acids (a. a.) and a MW of 70 kDa. We identified two proteoforms of its C-terminal fragments: PFR 8996840 (a.a. 528-622, 10.9 kDa) and PFR 8996850 (a.a. 515-622, 12.2 kDa). Another example is Rho GTPase-activating protein 45 (accession #Q92619-1), a 1136-a.a. protein with a 125 kDa MW. Its N-terminal fragment (PFR 9014676, a.a. 1-69, 7.2 kDa) was found with N-terminal acetylation. These two examples illustrate the importance of using TDP to acquire proteoform level information to accurately assign protein fragments and PTM sites and associate with their biological functions, which may be drastically different from their canonical form and will be neglected and indistinguishable by simply assigning them into a protein group in BUP approach.

### Detection of Low-Abundant Proteins and Their Proteoforms

Next, we investigated the depth of proteome coverage afforded via NP-based plasma enrichment. We mapped the proteins identified in each treatment condition (neat plasma, NPs) to the HPPP database for their rankings and estimated concentrations in plasma to understand whether NPs enrich low-abundant proteins.^31^ The results are illustrated in waterfall plots with x-axis of concentration ranking and y-axis of estimated concentration (Figure 3A-C, left). Each circle represents a protein identified in this study with TDP and the five least abundant proteins in each condition are labeled with their UniProt accession numbers. Figure 3A shows that in neat plasma, the proteoform identification was limited to proteins that had an estimated concentration above 1,000 ng/mL from the top 10% population in plasma proteome. While in Figure 3B and **3C**, the identification was made significantly deeper with NP enrichment enabling detection of low-abundant proteins. Remarkably, the least abundant protein that we identified was V-type proton ATPase 16 kDa proteolipid subunit (accession #P27449), which had an estimated concentration of 9 pg/mL and ranked 4027 by HPPP (**Table S1**).^31^ The proteoform of this ATPase subunit (PFR 1499) was found with N-terminal methionine removal and acetylation.

**Figure 3.**
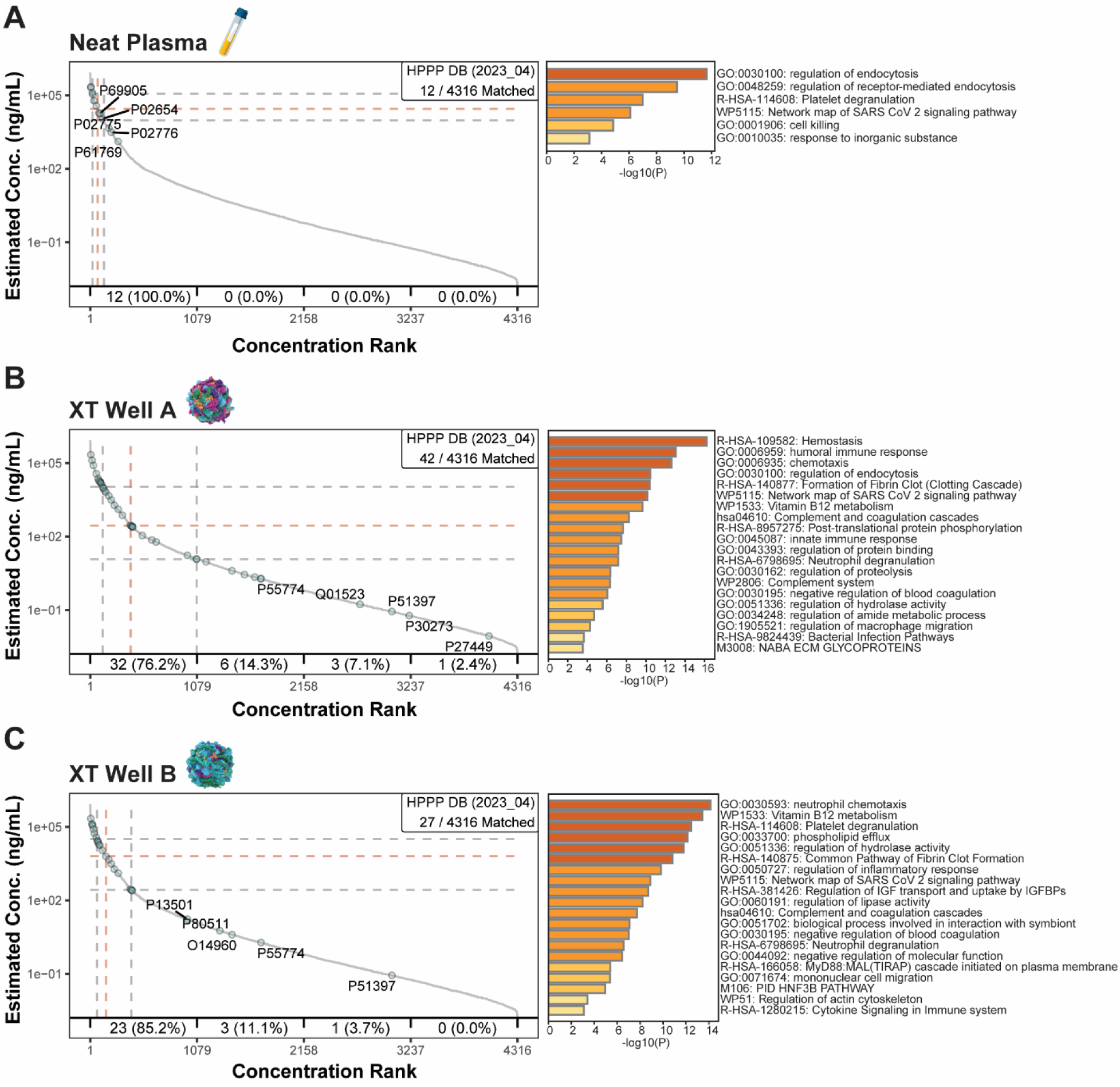
Waterfall plots of identified proteins and their abundances reported in HPPP (left) and gene ontology analysis (right). The individual waterfall plots of protein identified in neat plasma (A) and XT NPs (B-C) show that the NPs enable the detection of more proteoforms of proteins that have low estimated abundances in the HPPP. GO analysis revealed that more proteins associated with immune responses and cell signaling were identified in NP-enriched conditions.

Furthermore, the HPPP concentration estimation is based on protein level information and does not factor in signal dilution by complicated proteoform landscapes of each protein. For example, we previously found that ApoA-I, despite being the 11^th^ most abundant protein in plasma, had a wide dynamic range within its own proteoform landscape. Those comprised of less than 5% (glyco-proteoforms) and <2% (acylated proteoforms) of total ApoA-I were far more correlated with indices associated with risk of cardiovascular disease.^26^ Therefore, proteoform-specific enrichment is also of particular interest in TDP analysis that distinguishes it from BUP. An example in this work is the identification of death-associated protein 1 (DAP1, accession #P51397) and its phospho-proteoforms (Figure 4A). DAP1 is a low-abundant plasma protein that has an estimated concentration of 0.09 ng/mL and ranks 3047 in the HPPP database (**Table S1**).^31^ It was not identified in neat plasma but was found in both XT well A and B after NP enrichment (see waterfall plots in Figure 3B and **3C**). Interestingly, in XT well B, only the canonical proteoform (PFR 2628. a.a. 2-102, with N-methionine removal and acetylation, Figure 4B) was identified. However, in XT well A, there were four additional proteoforms (PFR 8616, 13512, 15907 and 9039157) identified with various combinations of PTMs at different sites including phosphorylations at S48 and S50, acetylation at K28 and N-terminal truncation from cleavage between G21 and G22 (Figure 4C, full proteoform sequence can be found in **Table S2**). The identifications described here are based on a 1% FDR cutoff. To further validate them, we used label-free quantification (LFQ) to compare proteoform abundances in each treatment condition (see box plots in **Figure S1A**). We found that the intensity of the canonical DAP1 proteoform (PFR 2628) was significantly higher in XT well A and B than in neat plasma, supporting that it was identified in both NP wells, and the DAP1 phospho-proteoforms (PFR 8616, 13512, and 9039157) were all significantly more abundant in NP well A compared to well B indicating differential enrichment. Considering the diversity of the proteoform landscape of DAP1, each proteoform could be well under the estimated concentration of 0.09 ng/mL. The differential enrichment of phosphorylated DAP1 proteoforms between XT well A and B can be utilized for understanding proteoform specific changes in signaling.

**Figure 4.**
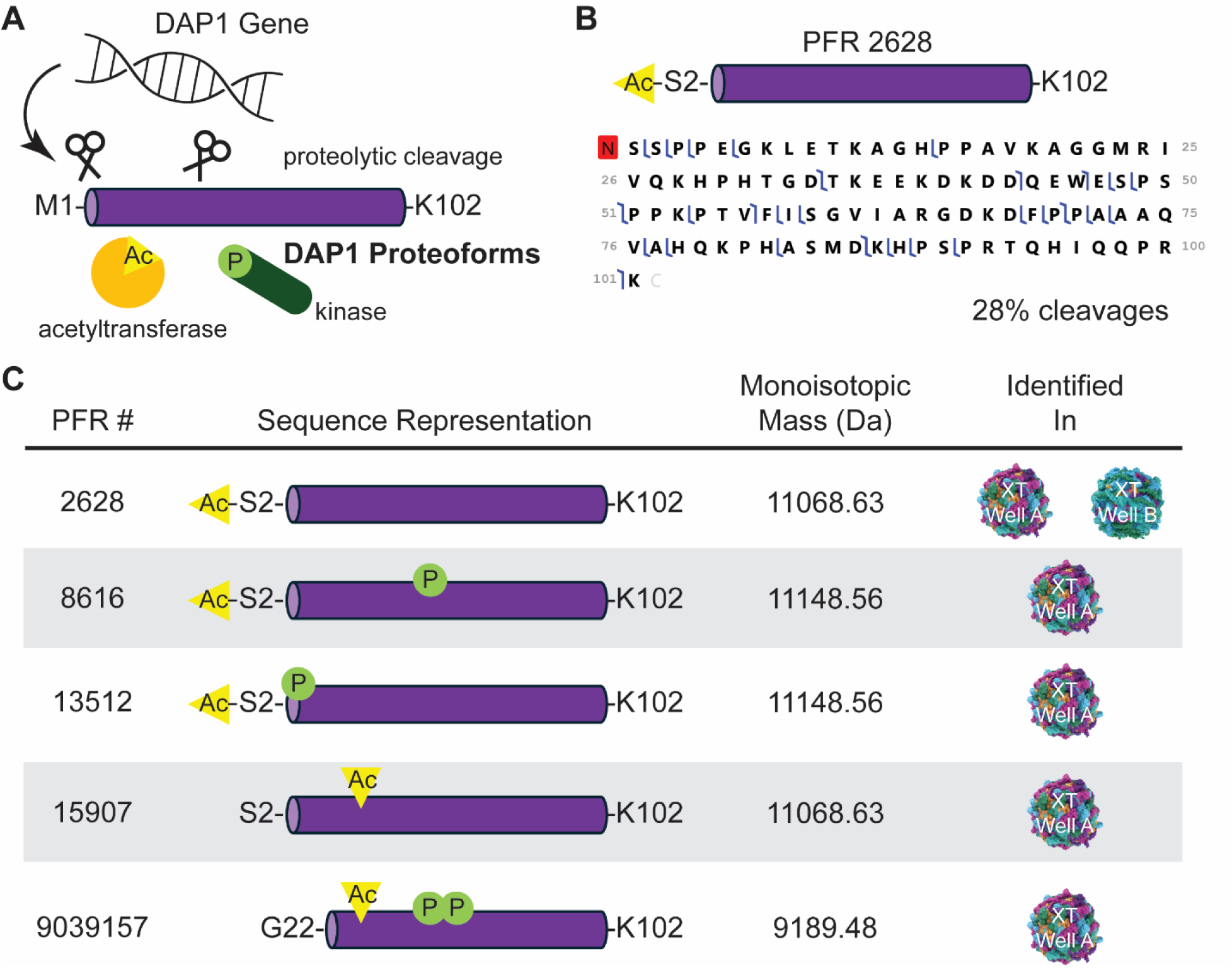
DAP1 proteoforms. (A) DAP1 protein undergoes several post-translation modifications including proteolytic cleavage, acetylation and phosphorylation that result in several proteoforms. (B) Representative sequence coverage map of one DAP1 proteoform (PFR 2628). (C) Five DAP1 proteoforms were identified in NP-enriched conditions.

The depth of the plasma proteome is connected with myriad molecular functions. For example, proteins secreted from tissues that regulate cell adhesion, signaling, and developmental functions as well as cytokines associated with immunity are ranked >1000 in concentration.^39–41^ We performed gene ontology (GO) analysis using all the protein accession numbers identified in each condition. The results are displayed on the right-hand side of each waterfall plot in Figure 3.

The proteins identified in neat plasma are limited to endocytosis, blood coagulation and acute responses, consistent with the function of the most abundant proteins in plasma. In XT well A and B, the proteome depth identified increases drastically even beyond the top 1000 proteins, revealing pathways associated with immune response and cell signaling. For example, we identified several low abundant chemokines including C-C motif chemokines 5, 14, and 18 (accession #P13501, #Q16627 and #P55774, respectively). The majority of C-C motif chemokine 5 proteoform landscape was comprised of PFR 40350, a fragment (a.a. 26-91, 7.6 kDa) of the canonical sequence found in both XT well A and B (**Table S3**). Additionally in XT well A, we found two less abundant proteoforms: PFR 18966 (a.a. 24-91, 7.8 kDa) with two additional residues at the N-terminus and PFR 5015352 (a.a. 26-91, 7.6 kDa) having the same length as PFR 40350 but with disulfide bond between C34 and C57. Similarly for C-C motif chemokine 18 (**Table S3**), we identified two proteoforms, PFR 53607 (a.a. 21-89. 7.9 kDa) and PFR 155657 (a.a. 21-88. 7.8 kDa), in both XT well A and B containing single amino acid variants (SAAVs) as well as an intra-molecular disulfide bond between C30 and C54 in PFR 5019691 (a.a. 21-89. 7.9 kDa). The LFQ box plots of C-C motif chemokine 5 and 18 proteoforms can be found in **Figure S1B** and **S1D**. The ability to identify and quantify these chemokines and their disulfides at the proteoform level is important for understanding their function and associated immunoresponses. For example, the disulfide between C30 and C54 in C-C motif chemokine 18 is known as essential for its activity.^42^ We found that in healthy donors, the ratio of reduced vs disulfide proteoforms (PFR 53607 vs 5019691) is roughly 1.25:1 (**Table S3**). It will be relevant to correlate the activation of the chemokine with diseased patients to understand their immune responses. We also note that the presence of disulfides would be lost in the routine BUP workflow in the reduction-alkylation step.

Lastly, we identified two C-C motif chemokine 14 proteoforms: PFR 98045 (a.a. 20-93. 8.7 kDa) and PFR 98050 (a.a. 20-93. 8.7 kDa, K61E). Instead of a PTM, we found a natural variant (K61E) that was reported in the dbSNP database (ref: rs16971802).^43^ Both C-C motif chemokine 14 proteoforms were only identified in XT well B with the 1% FDR threshold (**Table S3**). Curiously, the LFQ data showed contradicting results where there were no significant differences (PFR 98045) or little significance due to large variances in XT well B (PFR 98050) between neat plasma and NP enrichment conditions (**Figure S1C**). We speculated that this variance resulted from individuals expressing the different variants and sought to compare wild type (WT, PFR 98045) and K61E (PFR 98050) proteoform expression in each plasma donor. Box plots in **Figure S2** show that WT was differentially expressed in all three donors and K61E was detected at the level of noise in biorep01. The fold-changes of WT/K61E were 54, 0.24 and 0.1 for biorep01, biorep02 and biorep03 respectively with all *p*-values <2e-5. The potential difference in ionization efficiency due to the single site mutation was not considered in this analysis. However, the drastic fold-changes in the LFQ data still pointed to the conclusion where the donor for biorep01 only expressed WT, whereas the donor for biorep03 was homozygous for the K61E variant and therefore expressed no WT proteoform. Biorep02 had a slightly significant difference (*q* = 0.03) in WT expression compared to biorep03 so we could not rule out that the donor for biorep02 co-expressed both proteoforms. However, the ratio of WT/K61E in the biorep02 was 0.24, different from the 1:1 ratio that one would expect from a heterozygous gene variation (i.e. one copy of each in the genome). Therefore, further genome sequencing data of this individual will be complimentary to TDP to address whether the single-site mutation contributes to the proteoform stability, resistance to degradation, or upregulation in gene expression. It is also unknown how widespread the variant is and whether it is functionally different. This example demonstrates that TDP quantitatively captures nuanced differences in low-abundant chemokine proteoforms that may assist the development of precision medicine in immunology. In addition to genomics, we now have a new tool to study gene variants at the proteoform level, which tightly connects their composition to their molecular functions.

Our group has deep interests in studying proteoforms in blood to identify rejection biomarkers for organ transplantation.^21–25^ These studies were mostly performed using PBMCs, which is less accessible, and less uniform compared to plasma samples. The NP-enrichment allows us to quantify proteoforms involved in immune responses as well as several proteoform we previously identified as “immunoproteoforms” in a panel that could be potential biomarkers for liver-transplant outcome such as PFR 18628, 18631 (platelet factor 4, accession #P02776) and PFR 1464 (thymosin beta-4, accession #P62328).^25^ This workflow will enable the study of immunoproteoform responses in plasma with greater depth via TDP to expand the impacts of translational proteomics in organ transplant and autoimmune disease research.

### Nanoparticles Reproducibly Fractionate Proteoforms in Plasma

To further demonstrate that NPs differentially fractionate proteoforms from neat plasma and from each other, we took a quantitative TDP approach. The volcano plots comparing neat plasma versus XT well A (Figure 5A) and neat plasma versus XT well B (Figure 5B) clearly reflect the differences of proteoforms identified in neat plasma and NP-enriched conditions. The proteoforms on the right side of the plots in red indicate enrichment in neat plasma and the left sides in blue indicate enrichments in NPs. We observed differential proteoform fractionation of over 1024 (2^10^)-fold differences in some cases, indicating an extreme differential and reproducible affinity for NPs which were not observed in neat plasma. Figure 5C compares the two NP-enrichment conditions we used─ XT well A versus XT well B. The high confidence in large fold changes (>2^5^-fold) again supports that the two wells were engineered to interrogate largely orthogonal proteins and their proteoforms with little overlap. The use of both wells improves the proteome coverage and serves as another level of plasma protein fractionation─ not by MW but their affinities to the functionality of engineered NPs to form unique protein coronas.^11^

**Figure 5.**
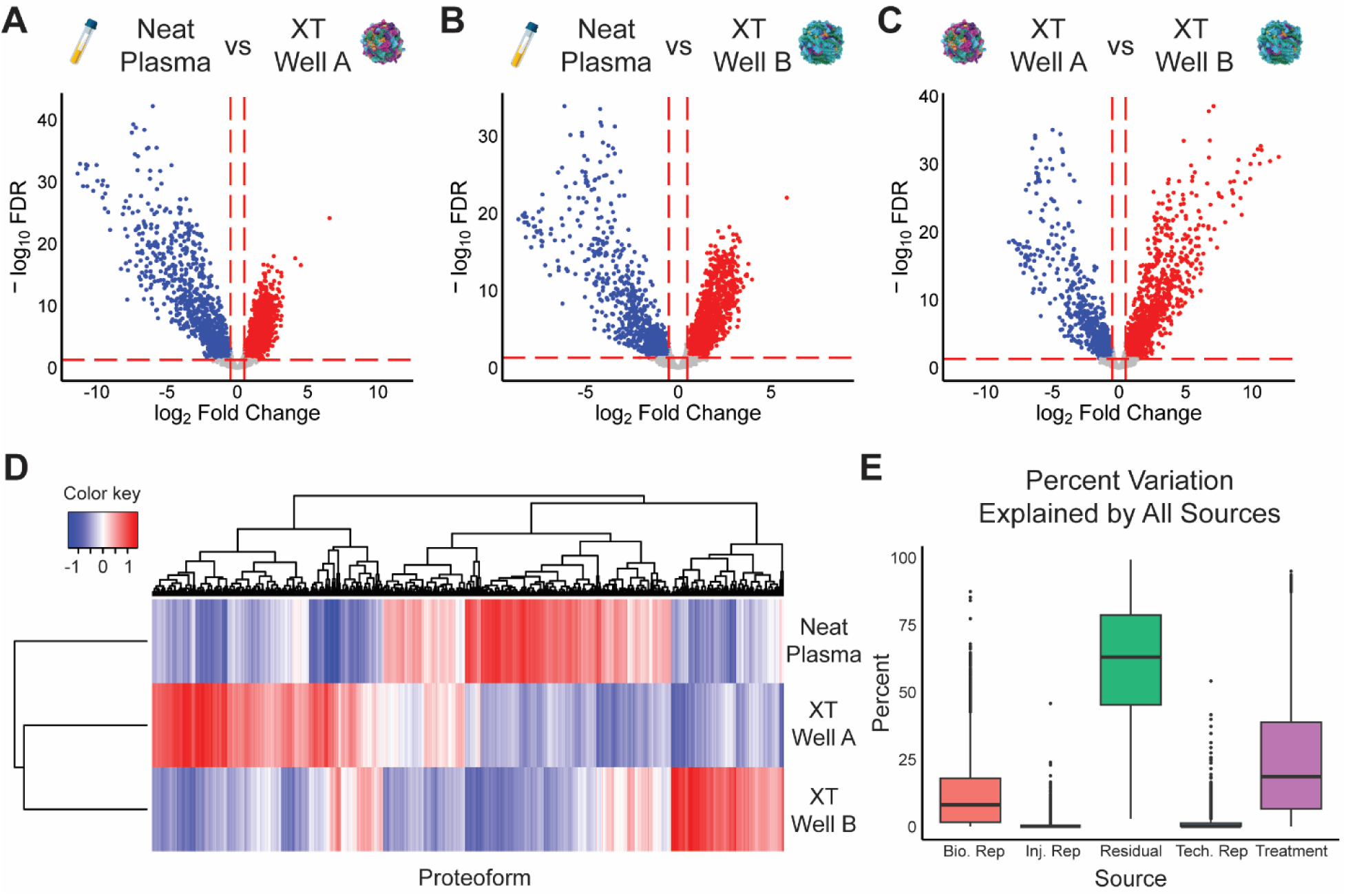
Nanoparticles differentially fractionate proteoforms in human plasma. (A-C) Volcano plots comparing neat plasma/NP and different NPs. Differential enrichments were characterized by large fold changes in proteoform abundances between neat plasma and nanoparticles (A-B) as well as between different nanoparticles (C). (D) Unsupervised clustering reveals unique clusters of proteoforms in neat plasma and enriched by different NPs. (E) Source of variance plot shows that the major contributor of differences observed are from different NP treatments.

To assess the effect of NP enrichment in another way, we next performed unsupervised clustering of proteoform abundances by three treatment conditions (neat plasma, XT well A, and XT well B). In the heatmap output (Figure 5D), each condition is represented with a unique cluster of enriched proteoforms, resulting in three distinct proteoform profiles. Sample clustering suggests that both NP-enriched samples are more similar to each other than the neat plasma, consistent with the bigger proteoform and protein intersections observed in Figure 2A and **2B**. Furthermore, the enriched clusters represent proteoforms uniquely identified in each treatment condition, correlating with additional GO functions revealed by NPs in Figure 3. In summary, these quantitative analyses support that NPs differentially interrogate proteoforms in plasma.

A box plot of percent variation explained by all sources (Figure 5E) including biological replicates (different plasma donors), technical replicates (complete sample preparation procedure including Proteograph NP-enrichment and PEPPI steps), injection replicates (multiple LC-MS injections of the same samples), treatments (neat plasma, XT well A and XT well B) and residual (all other potential sources) shows that the majority of variance outside of residual came from the different NP treatments. This is consistent with the observation that NPs differentially interrogate proteoforms described above. The next largest variation source came from biological reps, representing differences in proteoforms from individuals. In **Figure S3**, we compare proteoforms from different donors in different treatment conditions (neat plasma, NPs) using volcano plots. We found that both neat plasma and two NP-enrichment wells allowed quantitative comparison of proteoform abundances between individuals. We note that the fold-changes (<2^5^) and confidence level (-log_10_FDR <15) were much smaller than the comparison between treatments in Figure 5A-C, consistent with the smaller variance explained in Figure 5E. We also observed that biorep01 may be more different from biorep02 and biorep03 from more significantly changing proteoforms in volcano plots in all three treatment conditions (**Figure S3**), including the C-C motif chemokine 14 gene variation discussed above and highlighted in **Figure S2**. While the sample size and additional information about the donors are limited, the observation points to future application of the assay in analyzing clinical samples from patients for precision medicine. Lastly, both technical and injection replicates demonstrated low levels of variation, suggesting that NP-enrichment using the Proteograph workflow combined with PEPPI-MS for TDP is robust and reproducible.

### Limitation and Future Directions

While we report a new record number of proteoforms identified and depth in plasma, this work is still limited by common challenges in TDP, particularly in the analysis of high MW proteoforms and the amount of material needed. As described here, we began with an SDS-compliant workflow to assess the potential and effect size of the NP approach but the analysis of large proteoforms (>50 kDa) by LC-MS remains challenging due to several restrictions: 1) Instrument resolving power and ion decay─ limiting acquisition of isotopically-resolved spectra of large/highly charged proteoform ions.^37^ 2) Separation technology─ proteoforms are poorly separated due to peak broadening and overlapping, resulting in extremely complex spectra that are hard to deconvolute. 3) Inefficient sample preparation and clean-up─ due to the challenge in separation, fractionation of samples is often needed. In addition, these methods require precipitation of intact proteins to remove detergents, which typically requires >100 µg of total proteins.^24, 32^ Development of a streamlined protocol to deliver protein corona to an efficient TDP process will greatly enable this approach.

New acquisition methods continue to improve detection of large proteoforms. For examples, Ge and co-workers recently showed that using LC-MS combined with charge state deconvolution (low resolution, not isotopically resolved) can detect proteoforms up to 220 kDa for highly abundant proteins in specific tissues.^44^ Sun and co-workers combined CZE separation and low resolution MS^1^ acquisition to achieve plasma proteoform characterization in 3-70 kDa “mid” mass range.^30^ However, low injection volumes (nL scale) substantially limit the sensitivity of CZE-MS and its ability to detect low abundant proteoforms.^45^ Although limited to <30 kDa proteoforms, Fornelli and co-workers’ unique FAIMS approach directly address the challenge by reducing the complexity of the spectra and minimizing the need for multiple fractionation steps in the TDP analysis of plasma and serum.^29^ Approaches such as proton transfer charge reduction (PTCR)^46^ and individual ion mass spectrometry (I^2^MS)^47, 48^ have recently been used for better MS^1^ and MS/MS analyses of large proteoforms and their fragments that can be applied to the study of plasma proteome.

Compared to BUP analyses of plasma proteome, this TDP workflow is still limited in proteome depth and the amount of material needed. Typically in BUP, one well of Proteograph XT enrichment is sufficient to characterize 1,500-2,000 proteins groups (**Figure S4**) using an Orbitrap Exploris 480 mass spectrometer while the current TDP workflow is limited by MW, abundance, and sample preparation efficiency that only allows the characterization of proteoforms from 114 proteins. Given that proteoforms are the new currency in proteomics,^20^ we are encouraged by the ability to detect proteoforms in the low pg/mL concentration range and continue to improve our sample preparation workflow. For example, we are experimenting with combining XT well A and B to improve protein recovery from gel fractionation as well as several detergent-free elution methods for direct infusion to LC-MS. As the proteomics field starts to recognize that proteoforms correlate more tightly to phenotypes in patient cohorts than tryptic peptides^18–20, 25^, continual improvement of sample preparation workflow and reducing the amount of sample needed will be crucial for the future application of plasma TDP in translational studies where patient samples are limited in quantity.

## CONCLUSIONS

Nanoparticle-based protein corona enrichment of low-abundant proteins is a fast-growing approach in deep proteomics profiling of plasma proteome. Here, we adapt the BUP-based Proteograph workflow for TDP analysis and report a record number of 2841 proteoforms identified from 114 plasma proteins, revealing pathways previously hindered by abundant proteins such as albumin and apolipoproteins. This approach will allow proteoform-level analysis and discovery of biomarkers in diseases, which are more closely related to molecular function than protein group-level information. The workflow can also be coupled with emerging TDP acquisition methods such as low-resolution MS^1^, CZE-MS, PTCR and I^2^MS to further reduce spectral complexity, improve separation and extend mass range beyond 50 kDa.

## ASSOCIATED CONTENT

### Supporting Information

- Figure S1, label-free quantification of selected low abundant proteoforms; Figure S2, two proteoforms of C-C motif chemokine 14 in different donors; Figure S3, comparison of samples from different donors; Figure S4, comparison of BUP and TDP using Proteograph XT NPs; Table S1, the five least abundant proteins in each treatment condition; Table S2, DAP1 proteoform; Table S3, chemokine proteoforms (PDF).
- Table S4, list of proteoforms (PFR #) and proteins (accession #) identified and proteoform intensity sheet (xlsx).

## Notes

M.B., G.H., A.S., X.Z., R.B., and A.S. are employees of Seer Inc., which has commercialized the Proteograph Product Suite. N.L.K. is involved in entrepreneurial activities in top-down proteomics and consults for Thermo Fisher Scientific. The other authors declare that they have no other competing interests.

## ACKNOWLEDGEMENTS

This research was supported by the National Resource for Translational and Developmental Proteomics [P41 GM108569].

## TABLE OF CONTENTS GRAPHICS

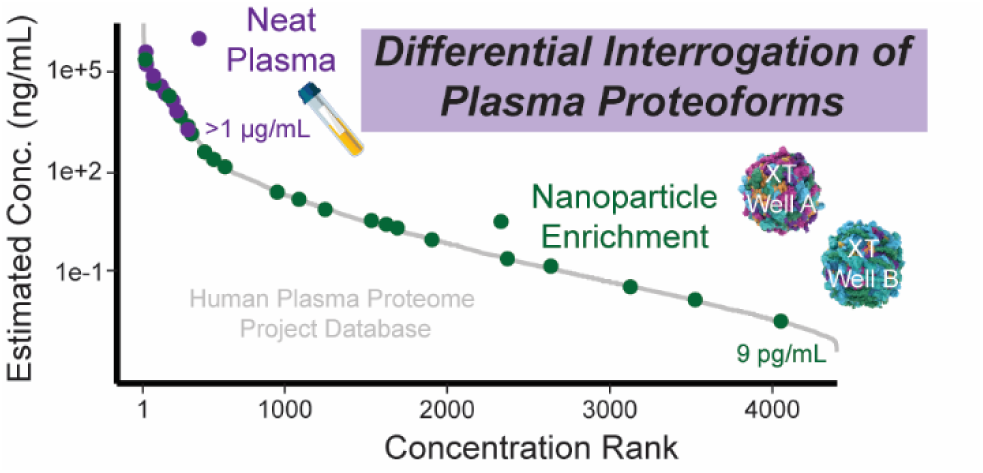

## Notes

https://doi.org/doi:10.25345/C53B5WK7J

## REFERENCES

(1) Surinova, S.; Schiess, R.; Huttenhain, R.; Cerciello, F.; Wollscheid, B.; Aebersold, R. On the development of plasma protein biomarkers. J. Proteome Res. 2011, 10 (1), 5–16. DOI: 10.1021/pr1008515.

(2) Geyer, P. E.; Holdt, L. M.; Teupser, D.; Mann, M. Revisiting biomarker discovery by plasma proteomics. Mol. Syst. Biol. 2017, 13 (9), 942. DOI: 10.15252/msb.20156297.

(3) Eldjarn, G. H.; Ferkingstad, E.; Lund, S. H.; Helgason, H.; Magnusson, O. T.; Gunnarsdottir, K.; Olafsdottir, T. A.; Halldorsson, B. V.; Olason, P. I.; Zink, F.;, et al. Large-scale plasma proteomics comparisons through genetics and disease associations. Nature 2023, 622 (7982), 348–358. DOI: 10.1038/s41586-023-06563-x.

(4) Hortin, G. L.; Sviridov, D. The dynamic range problem in the analysis of the plasma proteome. J. Proteomics 2010, 73 (3), 629–636. DOI: 10.1016/j.jprot.2009.07.001.

(5) Tirumalai, R. S.; Chan, K. C.; Prieto, D. A.; Issaq, H. J.; Conrads, T. P.; Veenstra, T. D. Characterization of the low molecular weight human serum proteome. Mol Cell Proteomics 2003, 2 (10), 1096–1103. DOI: 10.1074/mcp.M300031-MCP200.

(6) Shen, Y.; Kim, J.; Strittmatter, E. F.; Jacobs, J. M.; Camp, D. G., 2nd; Fang, R.; Tolie, N.; Moore, R. J.; Smith, R. D. Characterization of the human blood plasma proteome. Proteomics 2005, 5 (15), 4034–4045. DOI: 10.1002/pmic.200401246.

(7) Gianazza, E.; Miller, I.; Palazzolo, L.; Parravicini, C.; Eberini, I. With or without you - Proteomics with or without major plasma/serum proteins. J. Proteomics 2016, 140, 62–80. DOI: 10.1016/j.jprot.2016.04.002.

(8) Lee, P. Y.; Osman, J.; Low, T. Y.; Jamal, R. Plasma/serum proteomics: depletion strategies for reducing high-abundance proteins for biomarker discovery. Bioanalysis 2019, 11 (19), 1799–1812. DOI: 10.4155/bio-2019-0145.

(9) Fang, X.; Zhang, W.-W. Affinity separation and enrichment methods in proteomic analysis. J. Proteomics 2008, 71 (3), 284–303. DOI: 10.1016/j.jprot.2008.06.011.

(10) Bandow, J. E. Comparison of protein enrichment strategies for proteome analysis of plasma. Proteomics 2010, 10 (7), 1416–1425. DOI: 10.1002/pmic.200900431.

(11) Blume, J. E.; Manning, W. C.; Troiano, G.; Hornburg, D.; Figa, M.; Hesterberg, L.; Platt, T. L.; Zhao, X.; Cuaresma, R. A.; Everley, P. A.;, et al. Rapid, deep and precise profiling of the plasma proteome with multi-nanoparticle protein corona. Nature Communications 2020, 11 (1), 3662. DOI: 10.1038/s41467-020-17033-7.

(12) Ferdosi, S.; Stukalov, A.; Hasan, M.; Tangeysh, B.; Brown, T. R.; Wang, T.; Elgierari, E. M.; Zhao, X.; Huang, Y.; Alavi, A.;, et al. Enhanced Competition at the Nano–Bio Interface Enables Comprehensive Characterization of Protein Corona Dynamics and Deep Coverage of Proteomes. Adv. Mater. 2022, 34 (44), 2206008. DOI: 10.1002/adma.202206008.

(13) Ferdosi, S.; Tangeysh, B.; Brown, T. R.; Everley, P. A.; Figa, M.; McLean, M.; Elgierari, E. M.; Zhao, X.; Garcia, V. J.; Wang, T.;, et al. Engineered nanoparticles enable deep proteomics studies at scale by leveraging tunable nano–bio interactions. Proceedings of the National Academy of Sciences 2022, 119 (11), e2106053119. DOI: 10.1073/pnas.2106053119.

(14) Koh, B.; Liu, M.; Almonte, R.; Ariad, D.; Bundalian, G.; Chan, J.; Choi, J.; Chou, W.-F.; Cuaresma, R.; Hoedt, E.;, et al. Multi-omics profiling with untargeted proteomics for blood-based early detection of lung cancer. medRxiv 2024, 2024.2001.2003.24300798. DOI: 10.1101/2024.01.03.24300798.

(15) Donovan, M. K. R.; Huang, Y.; Blume, J. E.; Wang, J.; Hornburg, D.; Ferdosi, S.; Mohtashemi, I.; Kim, S.; Ko, M.; Benz, R. W.;, et al. Functionally distinct BMP1 isoforms show an opposite pattern of abundance in plasma from non-small cell lung cancer subjects and controls. PLoS One 2023, 18 (3), e0282821. DOI: 10.1371/journal.pone.0282821.

(16) Hendricks, N. G.; Bhosale, S. D.; Keoseyan, A. J.; Ortiz, J.; Stotland, A.; Seyedmohammad, S.; Nguyen, C. D. L.; Bui, J.; Moradian, A.; Mockus, S. M.;, et al. An inflection point in high-throughput proteomics with Orbitrap Astral: analysis of biofluids, cells, and tissues. bioRxiv 2024, 2024.2004.2026.591396. DOI: 10.1101/2024.04.26.591396.

(17) Vitko, D.; Chou, W.-F.; Nouri Golmaei, S.; Lee, J.-Y.; Belthangady, C.; Blume, J.; Chan, J. K.; Flores-Campuzano, G.; Hu, Y.; Liu, M.;, et al. timsTOF HT Improves Protein Identification and Quantitative Reproducibility for Deep Unbiased Plasma Protein Biomarker Discovery. J. Proteome Res. 2024, 23 (3), 929–938. DOI: 10.1021/acs.jproteome.3c00646.

(18) Smith, L. M.; Kelleher, N. L. Proteoform: a single term describing protein complexity. Nat. Methods 2013, 10 (3), 186–187. DOI: 10.1038/nmeth.2369.

(19) Aebersold, R.; Agar, J. N.; Amster, I. J.; Baker, M. S.; Bertozzi, C. R.; Boja, E. S.; Costello, C. E.; Cravatt, B. F.; Fenselau, C.; Garcia, B. A.;, et al. How many human proteoforms are there? Nat. Chem. Biol. 2018, 14 (3), 206–214. DOI: 10.1038/nchembio.2576.

(20) Smith, L. M.; Kelleher, N. L. Proteoforms as the next proteomics currency. Science 2018, 359 (6380), 1106–1107. DOI: 10.1126/science.aat1884.

(21) Savaryn, J. P.; Toby, T. K.; Catherman, A. D.; Fellers, R. T.; LeDuc, R. D.; Thomas, P. M.; Friedewald, J. J.; Salomon, D. R.; Abecassis, M. M.; Kelleher, N. L. Comparative top down proteomics of peripheral blood mononuclear cells from kidney transplant recipients with normal kidney biopsies or acute rejection. Proteomics 2016, 16 (14), 2048–2058. DOI: 10.1002/pmic.201600008.

(22) Toby, T. K.; Fornelli, L.; Srzentić, K.; DeHart, C. J.; Levitsky, J.; Friedewald, J.; Kelleher, N. L. A comprehensive pipeline for translational top-down proteomics from a single blood draw. Nat. Protoc. 2019, 14 (1), 119–152. DOI: 10.1038/s41596-018-0085-7.

(23) Huang, C.-F.; Su, P.; Fisher, T. D.; Levitsky, J.; Kelleher, N. L.; Forte, E. Mass spectrometry-based proteomics for advancing solid organ transplantation research. Frontiers in Transplantation 2023, 2, Mini Review. DOI: 10.3389/frtra.2023.1286881.

(24) Huang, C.-F.; Kline, J. T.; Negrão, F.; Robey, M. T.; Toby, T. K.; Durbin, K. R.; Fellers, R. T.; Friedewald, J. J.; Levitsky, J.; Abecassis, M. M. I.;, et al. Targeted Quantification of Proteoforms in Complex Samples by Proteoform Reaction Monitoring. Anal. Chem. 2024, 96 (8), 3578–3586. DOI: 10.1021/acs.analchem.3c05578.

(25) Melani, R. D.; Gerbasi, V. R.; Anderson, L. C.; Sikora, J. W.; Toby, T. K.; Hutton, J. E.; Butcher, D. S.; Negrao, F.; Seckler, H. S.; Srzentic, K.;, et al. The Blood Proteoform Atlas: A reference map of proteoforms in human hematopoietic cells. Science 2022, 375 (6579), 411–418. DOI: 10.1126/science.aaz5284.

(26) Wilkins, J. T.; Seckler, H. S.; Rink, J.; Compton, P. D.; Fornelli, L.; Thaxton, C. S.; Leduc, R.; Jacobs, D.; Doubleday, P. F.; Sniderman, A.;, et al. Spectrum of Apolipoprotein AI and Apolipoprotein AII Proteoforms and Their Associations With Indices of Cardiometabolic Health: The CARDIA Study. Journal of the American Heart Association 2021, 10 (17). DOI: 10.1161/jaha.120.019890.

(27) Forte, E.; Sanders, J. M.; Pla, I.; Kanchustambham, V. L.; Hollas, M. A. R.; Huang, C.-F.; Sanchez, A.; Peterson, K. N.; Melani, R. D.; Huang, A.;, et al. Top-Down Proteomics Identifies Plasma Proteoform Signatures of Liver Cirrhosis Progression. bioRxiv 2024, 2024.2006.2019.599662. DOI: 10.1101/2024.06.19.599662.

(28) Cheon, D. H.; Nam, E. J.; Park, K. H.; Woo, S. J.; Lee, H. J.; Kim, H. C.; Yang, E. G.; Lee, C.; Lee, J. E. Comprehensive Analysis of Low-Molecular-Weight Human Plasma Proteome Using Top-Down Mass Spectrometry. J. Proteome Res. 2016, 15 (1), 229–244. DOI: 10.1021/acs.jproteome.5b00773.

(29) Kline, J. T.; Belford, M. W.; Boeser, C. L.; Huguet, R.; Fellers, R. T.; Greer, J. B.; Greer, S. M.; Horn, D. M.; Durbin, K. R.; Dunyach, J.-J.;, et al. Orbitrap Mass Spectrometry and High-Field Asymmetric Waveform Ion Mobility Spectrometry (FAIMS) Enable the in-Depth Analysis of Human Serum Proteoforms. J. Proteome Res. 2023, 22 (11), 3418–3426. DOI: 10.1021/acs.jproteome.3c00488.

(30) Sadeghi, S. A.; Ashkarran, A. A.; Mahmoudi, M.; Sun, L. Mass spectrometry-based top-down proteomics in nanomedicine: proteoform-specific measurement of protein corona. bioRxiv 2024, 2024.2003.2022.586273. DOI: 10.1101/2024.03.22.586273.

(31) Deutsch, E. W.; Omenn, G. S.; Sun, Z.; Maes, M.; Pernemalm, M.; Palaniappan, K. K.; Letunica, N.; Vandenbrouck, Y.; Brun, V.; Tao, S.-c.;, et al. Advances and Utility of the Human Plasma Proteome. J. Proteome Res. 2021, 20 (12), 5241–5263. DOI: 10.1021/acs.jproteome.1c00657.

(32) Takemori, A.; Butcher, D. S.; Harman, V. M.; Brownridge, P.; Shima, K.; Higo, D.; Ishizaki, J.; Hasegawa, H.; Suzuki, J.; Yamashita, M.;, et al. PEPPI-MS: Polyacrylamide-Gel-Based Prefractionation for Analysis of Intact Proteoforms and Protein Complexes by Mass Spectrometry. J. Proteome Res. 2020, 19 (9), 3779–3791. DOI: 10.1021/acs.jproteome.0c00303.

(33) Giardine, B.; Riemer, C.; Hardison, R. C.; Burhans, R.; Elnitski, L.; Shah, P.; Zhang, Y.; Blankenberg, D.; Albert, I.; Taylor, J.;, et al. Galaxy: A platform for interactive large-scale genome analysis. Genome Res. 2005, 15 (10), 1451–1455.

(34) LeDuc, R. D.; Fellers, R. T.; Early, B. P.; Greer, J. B.; Shams, D. P.; Thomas, P. M.; Kelleher, N. L. Accurate Estimation of Context-Dependent False Discovery Rates in Top-Down Proteomics*[S]. Mol. Cell. Proteomics 2019, 18 (4), 796–805. DOI: 10.1074/mcp.RA118.000993.

(35) Ntai, I.; Toby, T. K.; LeDuc, R. D.; Kelleher, N. L. A Method for Label-Free, Differential Top-Down Proteomics. In Quantitative Proteomics by Mass Spectrometry, Sechi, S. Ed.; Springer New York, 2016; pp 121-133.

(36) Zhou, Y.; Zhou, B.; Pache, L.; Chang, M.; Khodabakhshi, A. H.; Tanaseichuk, O.; Benner, C.; Chanda, S. K. Metascape provides a biologist-oriented resource for the analysis of systems-level datasets. Nature Communications 2019, 10 (1), 1523. DOI: 10.1038/s41467-019-09234-6.

(37) Compton, P. D.; Zamdborg, L.; Thomas, P. M.; Kelleher, N. L. On the Scalability and Requirements of Whole Protein Mass Spectrometry. Anal. Chem. 2011, 83 (17), 6868–6874. DOI: 10.1021/ac2010795.

(38) Fornelli, L.; Toby, T. K. Characterization of large intact protein ions by mass spectrometry: What directions should we follow? Biochim. Biophys. Acta 2022, 1870 (4), 140758. DOI: 10.1016/j.bbapap.2022.140758.

(39) Tu, C.; Rudnick, P. A.; Martinez, M. Y.; Cheek, K. L.; Stein, S. E.; Slebos, R. J. C.; Liebler, D. C. Depletion of Abundant Plasma Proteins and Limitations of Plasma Proteomics. J. Proteome Res. 2010, 9 (10), 4982–4991. DOI: 10.1021/pr100646w.

(40) Geyer, Philipp E.; Kulak, Nils A.; Pichler, G.; Holdt, Lesca M.; Teupser, D.; Mann, M. Plasma Proteome Profiling to Assess Human Health and Disease. Cell Systems 2016, 2 (3), 185–195. DOI: 10.1016/j.cels.2016.02.015.

(41) Almeida, N.; Rodriguez, J.; Pla Parada, I.; Perez-Riverol, Y.; Woldmar, N.; Kim, Y.; Oskolas, H.; Betancourt, L.; Valdés, J. G.; Sahlin, K. B.;, et al. Mapping the Melanoma Plasma Proteome (MPP) Using Single-Shot Proteomics Interfaced with the WiMT Database. In Cancers (Basel), 2021; Vol. 13.

(42) Legendre, B.; Tokarski, C.; Chang, Y.; De Freitas Caires, N.; Lortat-Jacob, H.; Nadaï, P. D.; Rolando, C.; Duez, C.; Tsicopoulos, A.; Lassalle, P. The disulfide bond between cysteine 10 and cysteine 34 is required for CCL18 activity. Cytokine 2013, 64 (1), 463–470. DOI: 10.1016/j.cyto.2013.04.028.

(43) Sherry, S. T.; Ward, M. H.; Kholodov, M.; Baker, J.; Phan, L.; Smigielski, E. M.; Sirotkin, K. dbSNP: the NCBI database of genetic variation. Nucleic Acids Res. 2001, 29 (1), 308–311. DOI: 10.1093/nar/29.1.308.

(44) Melby, J. A.; Brown, K. A.; Gregorich, Z. R.; Roberts, D. S.; Chapman, E. A.; Ehlers, L. E.; Gao, Z.; Larson, E. J.; Jin, Y.; Lopez, J. R.; et al. High sensitivity top–down proteomics captures single muscle cell heterogeneity in large proteoforms. Proceedings of the National Academy of Sciences 2023, 120 (19), e2222081120. DOI: 10.1073/pnas.2222081120.

(45) Drown, B. S.; Jooß, K.; Melani, R. D.; Lloyd-Jones, C.; Camarillo, J. M.; Kelleher, N. L. Mapping the Proteoform Landscape of Five Human Tissues. J. Proteome Res. 2022, 21 (5), 1299–1310. DOI: 10.1021/acs.jproteome.2c00034.

(46) Huguet, R.; Mullen, C.; Srzentić, K.; Greer, J. B.; Fellers, R. T.; Zabrouskov, V.; Syka, J. E. P.; Kelleher, N. L.; Fornelli, L. Proton Transfer Charge Reduction Enables High-Throughput Top-Down Analysis of Large Proteoforms. Anal. Chem. 2019, 91 (24), 15732–15739. DOI: 10.1021/acs.analchem.9b03925.

(47) Kafader, J. O.; Melani, R. D.; Senko, M. W.; Makarov, A. A.; Kelleher, N. L.; Compton, P. D. Measurement of Individual Ions Sharply Increases the Resolution of Orbitrap Mass Spectra of Proteins. Anal. Chem. 2019, 91 (4), 2776–2783. DOI: 10.1021/acs.analchem.8b04519.

(48) McGee, J. P.; Melani, R. D.; Yip, P. F.; Senko, M. W.; Compton, P. D.; Kafader, J. O.; Kelleher, N. L. Isotopic Resolution of Protein Complexes up to 466 kDa Using Individual Ion Mass Spectrometry. Anal. Chem. 2021, 93 (5), 2723–2727. DOI: 10.1021/acs.analchem.0c03282.

