## Supplemental PDF for "Deep Profiling of Plasma Proteoforms with Engineered Nanoparticles for Top-down Proteomics"

**Table S4** .......... xlsx

### Figure S1

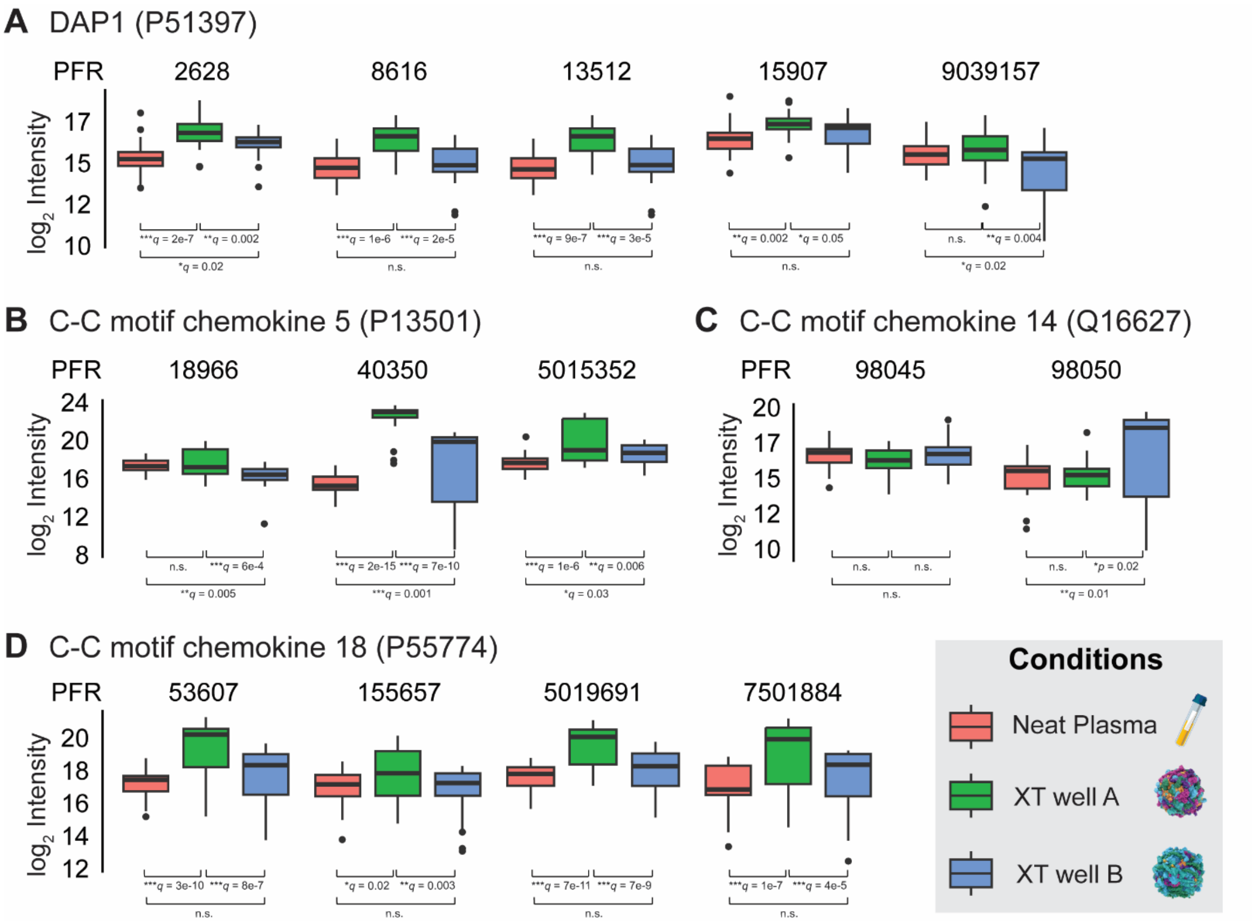

**Figure S1.** Label-free quantification of selected low abundant proteoforms of (A) DAP1, (B) C-C motif chemokine 5, (C) C-C motif chemokine 14, and (D) C-C motif chemokine 18. Significance codes: ***: q <0.001; **: q <0.01; *: q <0.05; n.s.: q >0.05.

#### Figure S2

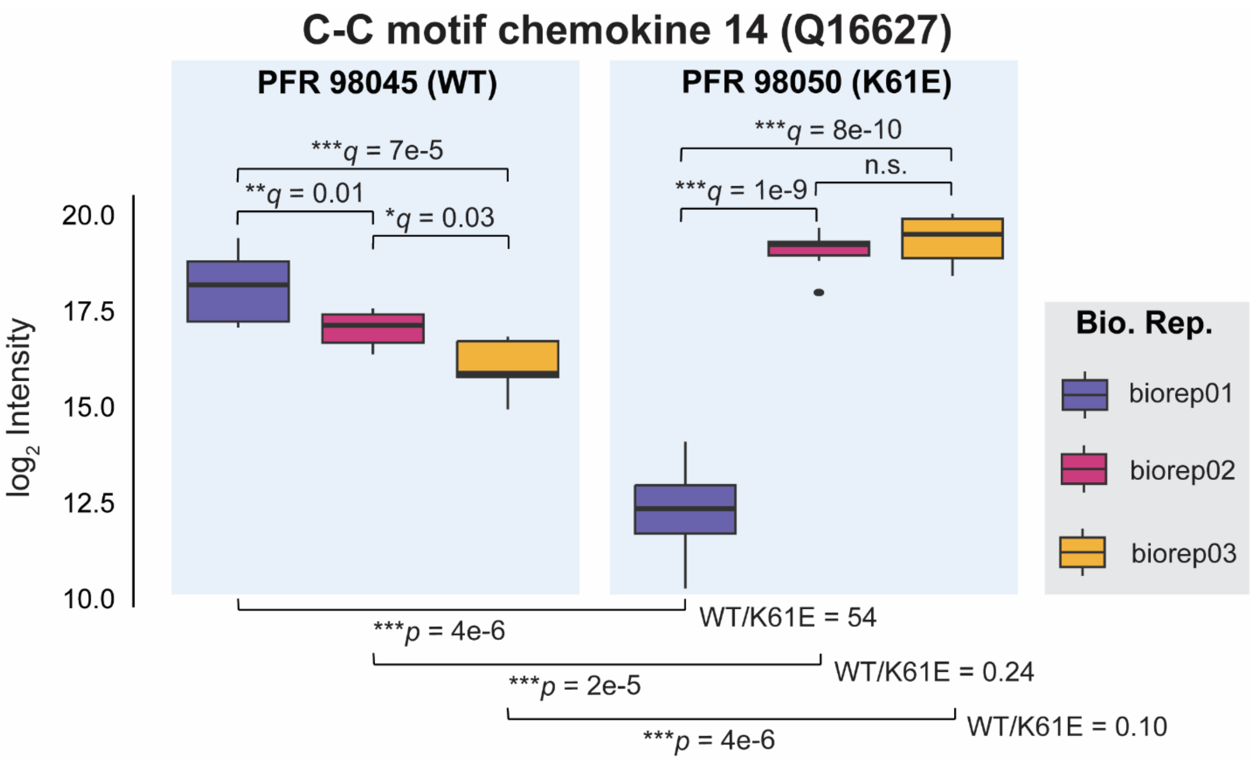

**Figure S2.** Two proteoforms of C-C motif chemokine 14 in different donors (XT well B only). Label-free quantification shows that biorep01 is significantly different from biorep02 and biorep03 in C-C motif chemokine 14 proteoform landscape. The drastic fold changes in WT/K61E in all three biological samples indicate that biorep01 expresses WT only (K61E proteoform at the level of noise) while biorep02 and biorep03 clearly express the K61E variant proteoform.

#### Figure S3

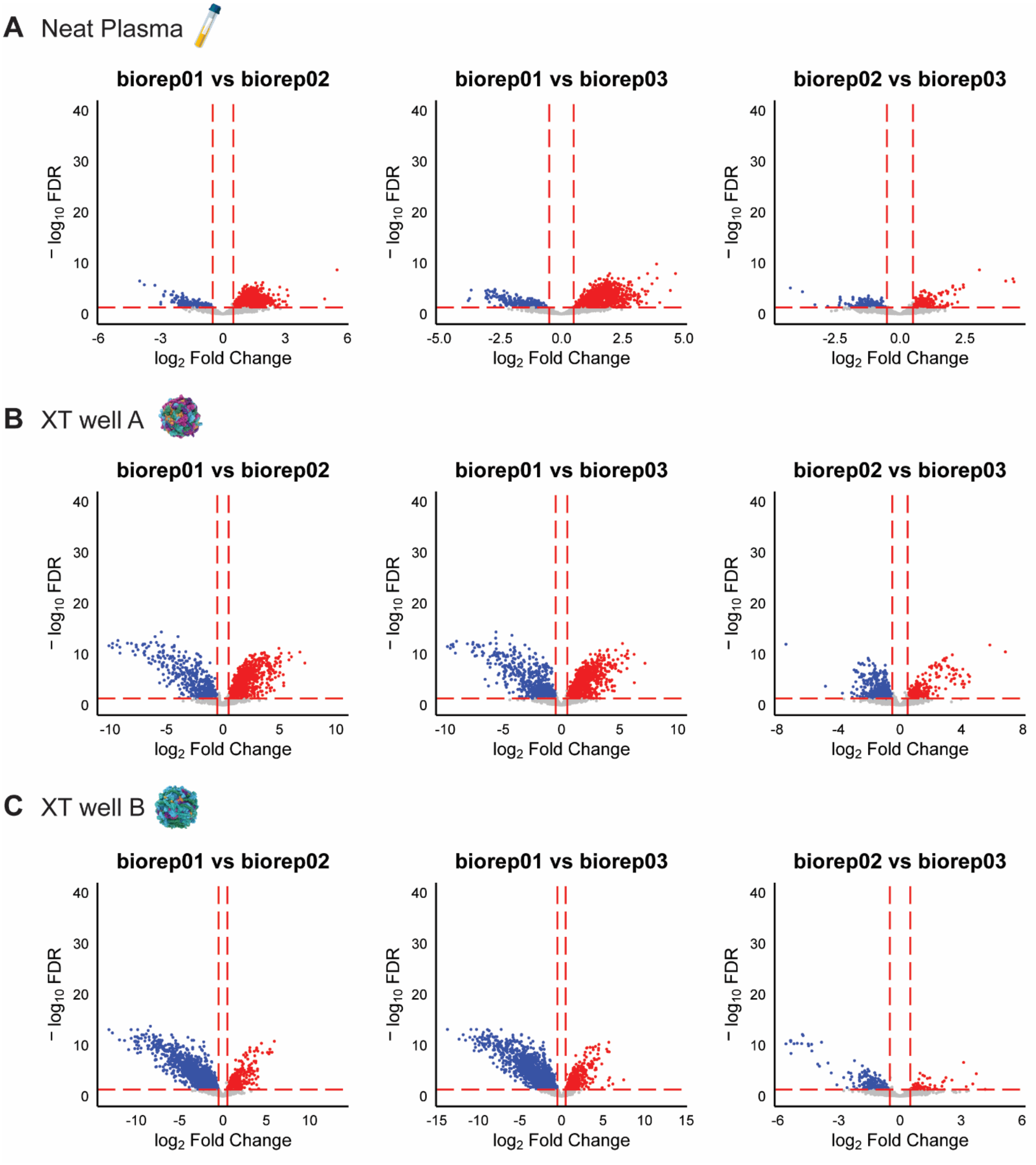

**Figure S3.** Volcano plot comparisons of proteoform abundances from the three donors in neat plasma (A), XT well A enrichment (B) and XT well B enrichment (C). Both fold-changes and confidence levels are much smaller compared to those between different treatments. Biorep01 seems to be more different than the other two donors.

#### Figure S4

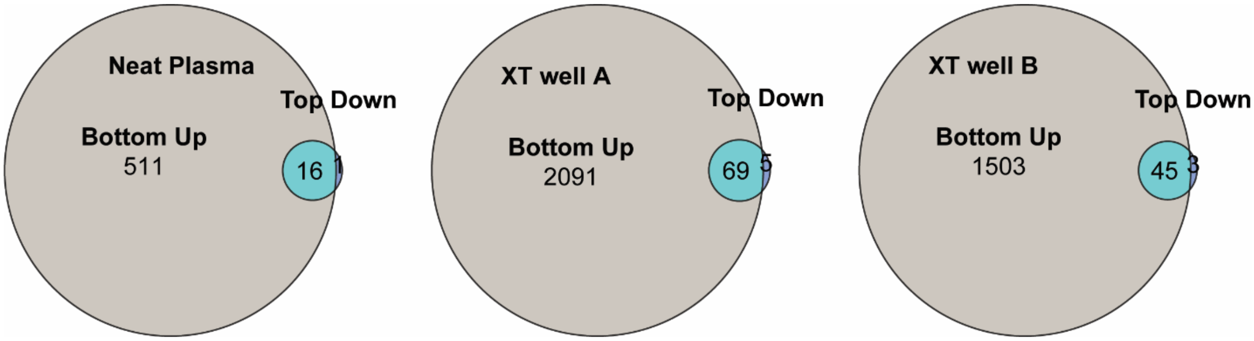

**Figure S4.** Comparing bottom-up and top-down protein identifications for neat plasma (left), XT well A enrichment (middle) and XT well B enrichment (right). For bottom-up analysis, samples from the same three donors were processed with standard Proteograph XT workflow and analyzed in Data-Independent Acquisition (DIA) mode with a Thermo Fisher Scientific Exploris 480 Mass spectrometer running a 30-min LC gradient. Raw MS data was processed using the DIA-NN search engine in library-free mode. Bottom-up results are reported as protein groups identified in all three technical replicates and union of biological replicates.

#### Table S1

**Table S1.** The 5 least abundant proteins (by HPPP estimates) identified in each treatment condition.

| Protein accession # | Treatment condition | HPPP estimated concentration (ng/mL) | Concentration ranking |
|---|---|---|---|
| **P27449** | XT Well A | 0.0087 | 4027 |
| **P30273** | XT Well A | 0.059 | 3222 |
| **P51397** | XT Well A,<br>XT Well B | 0.087 | 3047 |
| **Q01523** | XT Well A | 0.17 | 2725 |
| **P55774** | XT Well A,<br>XT Well B | 1.9 | 1725 |
| **O14960** | XT Well B | 4 | 1432 |
| **P80511** | XT Well B | 5.8 | 1309 |
| **P13501** | XT Well B | 17 | 982 |
| **P61769** | Neat | 1300 | 283 |
| **P02776** | Neat | 3100 | 213 |
| **P02775** | Neat | 4600 | 188 |
| **P02654** | Neat | 11000 | 123 |
| **P69905** | Neat | 18000 | 96 |

#### Table S2

**Table S2.** DAP1 (P51397) proteoforms.

| PFR | Sequence | Monoisotopic Mass (Da) | Identified In | Relative Abundance[a] |
|---|---|---|---|---|
| **2628** | Ac-SSPPEGKLETKAGHPPAVKAGGMRIVQK HPHTGDTKEEKDKDDQEWESPSPPKPT VFISGVIARGDKDFPPAAAQVAHQKP HASMDKHPSPRTQHIQQPRK | 11068.62932 | XT well A, XT well B | 2 |
| **8616** | Ac-SSPPEGKLETKAGHPPAVKAGGMRIVQK HPHTGDTKEEKDKDDQEWEpSPSPPKPT VFISGVIARGDKDFPPAAAQVAHQKP HASMDKHPSPRTQHIQQPRK | 11148.59565 | XT well A | 1 |
| **13512** | Ac-pSSPPEGKLETKAGHPPAVKAGGMRIVQK HPHTGDTKEEKDKDDQEWESPSPPKPT VFISGVIARGDKDFPPAAAQVAHQKP HASMDKHPSPRTQHIQQPRK | 11148.59565 | XT well A | 1 |
| **15907** | SSPPEGKLETKAGHPPAVKAGGMRIVQK(Ac) HPHTGDTKEEKDKDDQEWESPSPPKPT VFISGVIARGDKDFPPAAAQVAHQKP HASMDKHPSPRTQHIQQPRK | 11068.62932 | XT well A | 4 |
| **9039157** | GMRIVQK(Ac) HPHTGDTKEEKDKDDQEWEpSPpSPPKPT VFISGVIARGDKDFPPAAAQVAHQKP HASMDKHPSPRTQHIQQPRK | 9189.482052 | XT well A | 1.5 |

[a] Relative abundance was estimated from the raw intensities of the proteoforms in TDP LC-MS runs (MS[1]). The ionization efficiencies, matrix effects and other sources of bias were not considered.

#### Table S3

**Table S3.** Chemokine proteoforms.

| Protein | PFR | Sequence | Monoisotopic Mass (Da) | Identified In | Relative Abundance[a] |
|---|---|---|---|---|---|
| **C-C motif chemokine 5 (P13501)** | 18966 | SPYSSDTTPCCFAYIARPLPRA HIKEYFYTSGKCSNPAVVFVTRK NRQVCANPEKKWVREYINSLEMS | 7845.857 | XT well A | 1 |
| | 40350 | YSSDTTPCCFAYIARPLPRA HIKEYFYTSGKCSNPAVVFVTRK NRQVCANPEKKWVREYINSLEMS | 7661.772 | XT well A, XT well B | 50 |
| | 5015352 | YSSDTTPC(-H)CFAYIARPLPRA HIKEYFYTSGKC(-H)SNPAVVFVTRK NRQVCANPEKKWVREYINSLEMS | 7659.756 | XT well A | 1.5 |
| **C-C motif chemokine 14 (Q16627)** | 98045 | TKTESSSRGPYHPSECCFTYTTY KIPRQRIMDYYETNSQCSKPGIVFIT KRGHSVCTNPSDKWVQDYIKDMKEN | 8672.085 | XT well B | -[b] |
| | 98050 | TKTESSSRGPYHPSECCFTYTTY KIPRQRIMDYYETNSQCSEPGIVFIT KRGHSVCTNPSDKWVQDYIKDMKEN | 8673.033 | XT well B | -[b] |
| **C-C motif chemokine 18 (P55774)** | 53607 | AQVGTNKELCCLVYTSWQIPQKF IVDYSETSPQCPKPGVILLTKRGRQ ICADPNKKWVQKYISDLKLNA | 7850.098 | XT well A, XT well B | 2.5 |
| | 155657 | AQVGTNKELCCLVYTSWQIPQKF IVDYSETSPQCPKPGVILLTKRGRQ ICADPNKKWVQKYISDLKLN | 7779.061 | XT well A, XT well B | 1 |
| | 5019691 | AQVGTNKELC(-H)CLVYTSWQIPQKF IVDYSETSPQC(-H)PKPGVILLTKRGRQ ICADPNKKWVQKYISDLKLNA | 7848.082 | XT well A, XT well B | 2 |
| | 7501884 | AQVGTNKELC(-H)CLVYTSWQIPQKF IVDYSETSPQCPKPGVILLTKRGRQ ICADPNKKWVQKYISDLKLNA | 7849.09 | XT well A, XT well B | 2 |

[a] Relative abundance was estimated from the raw intensities of the proteoforms in TDP LC-MS runs ($MS^1$). The ionization efficiencies, matrix effects and other sources of bias were not considered.

[b] Varied in different donors, see **Figure S2**.
